# Exercising with virtual reality is potentially better for the working memory and positive mood than cycling alone

**DOI:** 10.1101/2024.05.07.593030

**Authors:** Genta Ochi, Ken Ohno, Ryuta Kuwamizu, Koya Yamashiro, Tomomi Fujimoto, Koyuki Ikarashi, Naoki Kodama, Hideaki Onishi, Daisuke Sato

**Author notes:** These authors contributed equally to this work (Genta Ochi, Ken Ohno). **Corresponding author:** Genta Ochi, 1398 Shimami-cho, Kita-ku, Niigata-City, 950-3198, Japan. **CRediT authorship contribution statement GO**: Funding acquisition, Conceptualization, Methodology, Data curation, Formal analysis, Writing – original draft, Writing – review & editing. **KO**: Funding acquisition, Methodology, Data curation, MRI analysis, Writing – original draft, Writing – review & editing. **RK**: Statistical analysis, Writing – review & editing. **KY**, **TF**, **KI, NK**: Writing – review & editing. **HO**: Funding acquisition, Writing – review & editing. **DS**: Funding acquisition, Conceptualization, Methodology, Writing – review & editing. **Funding** This work was supported in part by the Japan Society for the Promotion of Science (JSPS) Grant JP22K17739 (GO), JP23K14904 (KO), JP19H01090 (HO), 20K20621 (HO), and JP21H03310 (DS).

## Abstract

Although virtual reality (VR) exercise has attracted attention as a factor in exercise habituation due to its mood-enhancing effects, its impact on brain function remains unclear. This study, involving 23 healthy university students, used functional magnetic resonance imaging (fMRI) to explore how VR exercise affects working memory, a key executive function, and its underlying neural mechanisms. Our findings indicate that a 10-min VR exercise session improved mood (arousal and vitality level) and working memory task performance (3-back task) more effectively than exercise or rest alone. Furthermore, the results confirmed that increased vitality from exercise and VR exercise interventions was associated with improved 3-back task performance. However, specific brain regions contributing to this enhancement remain unidentified. These results highlight VR exercise as the optimal exercise program for enhancing working memory function by increasing vitality level. These insights underscore VR’s potential as a novel exercise modality with benefits extending beyond exercise adherence to potentially preventing dementia and depression.

## 1. Introduction

Previous studies have supported the benefits of physical activity in promoting physical and mental health. Furthermore, physical activity is becoming increasingly important as both older and younger adults have reported that high aerobic fitness is beneficial for maintaining executive function (Hillman, Erickson, and Kramer 2008; Weinstein et al. 2012; Kuwamizu et al. 2023; Hyodo et al. 2016). While transient exercise improves executive function (Yanagisawa et al. 2010; Byun et al. 2014; Kujach et al. 2018; Damrongthai et al. 2021), it does not always have this effect. Some studies report that exercise does not improve executive function (Yamazaki et al. 2018; Ishihara et al. 2021), indicating that factors beyond exercise intensity and duration may play a role in enhancing executive function.

A previous study revealed that exercising while listening to favorite music increased pleasant mood (arousal and pleasure level) and is related to the effect of exercise on improving executive function (Suwabe et al. 2021). Although the connection between pleasant mood and enhanced executive function is not well understood, the involvement of the brain’s catecholaminergic nervous system, which governs emotion, reward, and executive function, is conceivable. Utilizing the relationship between pupil diameter as an indirect measure of the activity of the locus coeruleus (LC) involved in psychological arousal (Yamazaki et al. 2023), Kuwamizu et al. (2023) suggested that the improvement in executive function induced by very light exercise is mediated by catecholaminergic neuron activity originating in the LC. This indicates that the activity of LC-derived catecholaminergic neurons is involved in the physiological condition for exercise-induced improvement of executive function.

Therefore, we focused on virtual reality (VR) as an environmental condition that elicits arousal (catecholaminergic neuronal activity) and pleasantness during exercise. VR environments are gaining attention as a new approach to promoting physical activity (Ahn and Fox 2017). Previous studies have suggested that VR may increase the potential for long-term participation in physical activity by distracting attention from negative images of exercise that depict it as physically fatiguing (Faric et al. 2019), boring, or strenuous (Qian, McDonough, and Gao 2020; Ekkekakis, Hall, and Petruzzello 2008). Furthermore, the authors have confirmed that VR exergames induce a pleasant mood (Ochi, Kuwamizu, Fujimoto, et al. 2022), and exercising in VR, which fosters positive mood, may enhance executive function (Byun et al. 2014; Suwabe et al. 2021; Fukuie et al. 2023). However, the clarity regarding whether exercise in VR enhances executive function compared to simple exercise or if LC activity (catecholaminergic neuron activity plays a role in the background, is currently lacking. Therefore, this study aimed to elucidate the effects of exercise in VR on executive function and its mechanisms in the brain using functional magnetic resonance imaging (fMRI).

The N-back task, a classic measure of working memory function, was chosen as the executive function task, and brain activity during the task was evaluated using fMRI to test whether exercise under VR enhances executive function via LC activity and by increasing activity in task-specific brain regions. In this task, participants monitor a series of stimuli and determine whether each presented stimulus matches the one presented N trials ago (N is a pre-specified integer, usually varying from 0–3). During the performance of this task, the stimuli are sequentially registered and memorized for a few seconds, and a motor response is required after each stimulus. The increased memory load in the N-back task presents a significant challenge, resulting in increased reaction time and a higher number of incorrect responses at the behavioral level. This task has been used in many studies examining the effects of exercise on executive function (Roig et al. 2013; Winter et al. 2007; McMorris et al. 2011; Pontifex et al. 2009; Weng et al. 2015; Gothe et al. 2013; Yamazaki et al. 2018). Furthermore, previous studies using fMRI have reported that the dorsolateral prefrontal cortex (DLPFC) is significant in the performance of N-back tasks and that DLPFC activity increases with increasing task difficulty (Lamichhane et al. 2020). Our previous study using the color-word Stroop task, in which DLPFC activity was considered as important as in the N-back task, confirmed that DLPFC activity increases when exercise enhances executive function (Yanagisawa et al. 2010; Byun et al. 2014; Hyodo et al. 2012; Kujach et al. 2018; Suwabe et al. 2021). Therefore, the hypothesis posits that improving executive function through VR exercise involves heightened DLPFC activity.

Therefore, in this study, we established three conditions: one where participants simply exercised, one where they rested, and another where they exercised in a VR environment using a head-mounted display (HMD). We aimed to determine whether VR exercise improves executive function more effectively than exercise and rest alone. This study suggests that VR enhances the effects of exercise on working memory.

## 2. Material and methods

### 2.1 Participants

Twenty-five healthy young Japanese adults with normal or corrected-to-normal vision participated in this study. We conducted a power analysis with Cohen’s *d* = 0.3 using behavior data from the executive task, referencing our previous study (Ochi et al. 2018; Ochi, Kuwamizu, Suwabe, et al. 2022). A power analysis using G-power (3.1.9.2; The G*Power Team) software showed that 24 subjects would be sufficient to detect a significant interaction in a repeated measure two-way analysis of variance (ANOVA) with 0.05 alpha and 80% power. All participants were right-handed and nonsmokers. No participant reported a history of respiratory, circulatory, or neurological disease or had an illness requiring medical care. All participants had normal or corrected-to-normal vision and normal color vision. Two participants lacked task proficiency, with correct response rates below 60%; therefore, data from the remaining 23 participants were used for the analysis. Post-hoc sensitivity analysis performed based on this sample with 80% power and α=.05 demonstrated sufficient sensitivity to detect interaction *f* = 0.306 as computed using G*Power. This study was conducted in accordance with the Declaration of Helsinki and approved by the appropriate ethics review board. Before participation, all participants were informed about the confidentiality of their data and provided written informed consent. Table 1 presents the demographic data of the participants.

**Table 1.** Participants’ characteristics.

| Characteristic | Female, n = 13<sup>1</sup> | Male, n = 10<sup>1</sup> | P-value<sup>2</sup> |
| --- | --- | --- | --- |
| Age (yr) | 19.5 (1.6) | 21.5 (2.5) | 0.04 |
| Height (cm) | 155.9 (4.8) | 172.2 (4.2) | <0.001 |
| Weight (kg) | 51.9 (8.4) | 69.5 (7.7) | <0.001 |
| Work rate (W) | 66.0 (16.2) | 96.0 (23.2) | 0.003 |
| $\dot{V}O_{2peak}$ (mL kg<sup>-1</sup> min<sup>-1</sup>) | 34.4 (6.6) | 40.4 (8.3) | 0.082 |

<sup>1</sup>Mean (SD).
<sup>2</sup>Welch two sample t-test: Differences between men and women were assessed.
Note: $\dot{V}O_{2peak}$ =peak oxygen uptake; SD=standard deviation.

### 2.2 Experimental procedures

On the first day, participants underwent a graded exercise test with a bicycle ergometer (Ergomedic 828E, MONARK, Sweden) to measure their peak oxygen uptake_•_ (*V*O_2peak_). The participants practiced the N-back task three times before being subjected to the main experimental conditions and engaged with the VR software. The participants were instructed to play an in-game tutorial to learn how the software is played. Once the tutorial was completed, the participants played the 10-min program once to familiarize themselves with VR.

A few days after the first visit, the participants engaged in one of three experimental conditions: exercise without HMD (EX), exercise with HMD (VR), or rest without HMD (RS)(Fig.1). Exercise sessions were conducted using the same bicycle ergometer setup as the graded exercise test. All participants completed all three conditions on a separate day, with the order counterbalanced across participants. In all conditions, the participants completed the N-back task before and immediately after 10 min of exercise using the MRI scanner. The participants completed a questionnaire before performing the N-back task outside the MRI scanner.

**Fig. 1.**
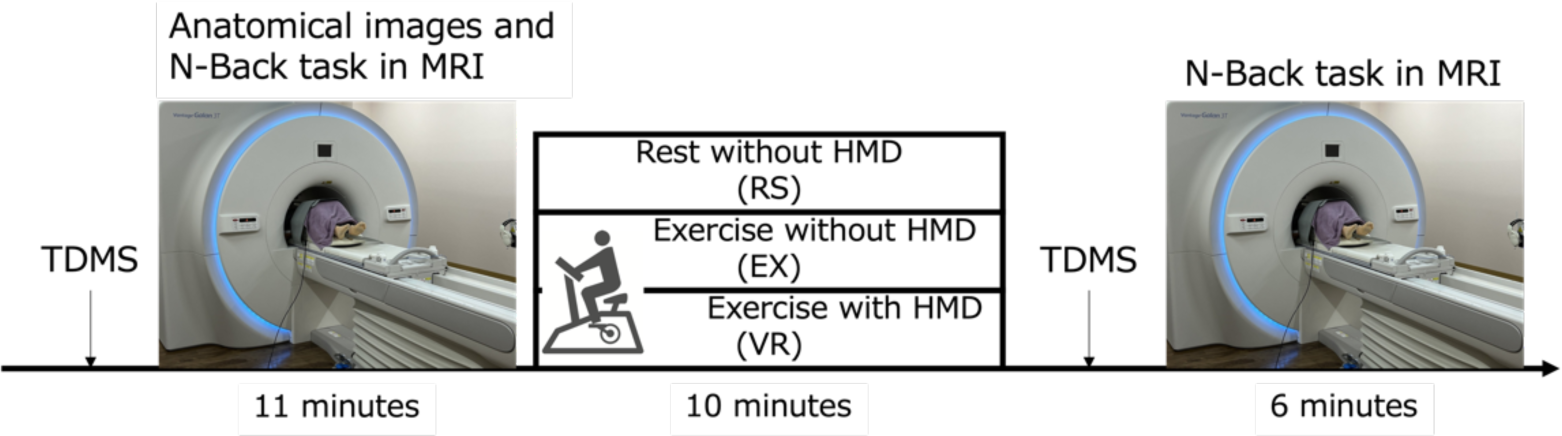
Experimental procedures.

TDMS, N-back task, and brain activity were measured before and after 10 min of exercise or rest. Before exercise, anatomical images were taken, followed by an N-back task and the fMRI images during the task. EX=exercise without HMD condition; VR=exercise with HMD condition; RS=rest without HMD condition; HMD=head-mounted display; TDMS=Two-Dimensional Mood Scale.

### 2.3 Virtual really environment

HOLOFIT (developed by Holodia AG) as the VR environment was exposed to participants using a commercially available HMD (Meta Quest 2, Meta Platforms, Inc.)(Fig. 2). HOLOFIT uses a cadence sensor (Wahoo Cadence Sensor, Wahoo Fitness), which causes the view to shift as the bicycle pedals rotate. In this study, all participants watched the Paris stage installed in HOLOFIT.

**Fig. 2.**
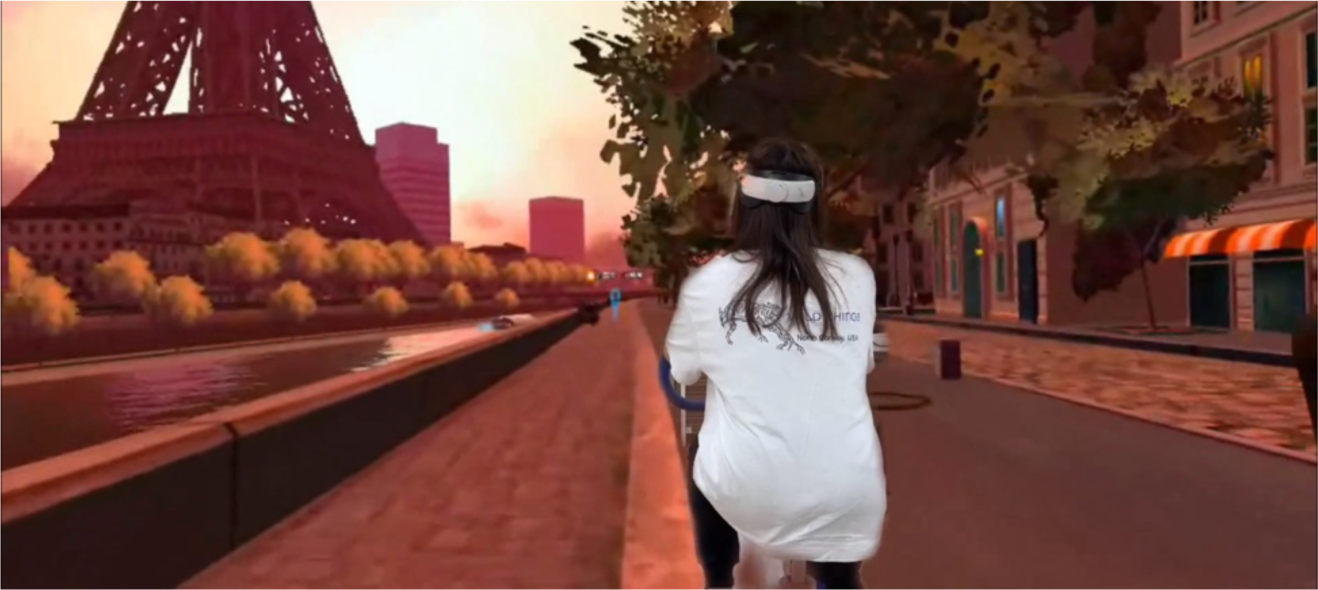
Images of the VR movement.

In the VR conditions, participants performed the bicycle exercise while wearing an HMD. This figure shows an image of a VR exercise that combines a participant on a bicycle and a format in which the participant is looking at the bicycle. VR=virtual reality; HMD=head-mounted display.

### 2.4 N-back task

Participants completed the color N-back tasks during the fMRI and outside the scanner. The color N-back task required participants to monitor a continuous color flow of single squares and respond when identical to the color presented at a specified interval (0, 1 or 3-back).

The N-back task paradigm consisted of three blocks of stimuli based on a previous study (Jacola et al. 2014). Each block contained 12 targets and 24 distractors in the 0 and 1, 3-back portions, respectively. Participants were informed of the target interval change via visually presented instructions: “red or not?” for 0-back, “Same as the one before?” for 1-back and “Every other two?” for 3-back. Stimuli were presented on a computer monitor for 0.5 s, with an inter-stimulus interval of 1.5 s. Each portion was measured 30 s apart.

Task performance yielded three outcome variables of interest: reaction time, number of omission errors (failure to respond to a target stimulus), and number of commission errors (response to a distractor stimulus). The reaction times were averaged across the load conditions for each participant. Performance accuracy was calculated separately for each modality and load condition using the following formula: Accuracy = Hits + Correct Rejections/Total Stimuli, where hits = number of targets – omission errors, and correct rejection = number of distractors – commission errors.

### 2.5 Cardiorespiratory aerobic fitness assessment

Individual aerobic fitness levels were determined using a graded exercise test with a bicycle ergometer (Ergomedic 828E, MONARK, Sweden) for determine the_•_ appropriate individual intensity for moderate exercise. *V* O_2peak_, the gold standard measurement of aerobic fitness, was determined by continuously measuring oxygen uptake during an incremental test to exhaustion. After warming up for 3 min at 30 W, the workload increased by 15 W·min^−1^ constantly and continuously until the maximal effort was reached. The pedal rotation speed was maintained at 60 rpm. Exhaled gas was analyzed using a gas analyzer (Aeromonitor AE-310S; Minato Medical Science, Osaka, Japan). The heart rate (HR) was measured during the assessment. The participants were asked to indicate their subjective feelings about exercise intensity using the Borg rating of perceived exertion (RPE) (15-point scale: 6 = no exertion; 20 = maximal exertion). All the participants exercised until they could no longer maintain a pedal rotation speed of 60_•_ rpm. *V*O_2peak_ was determined when at least two of the following criteria were satisfied: 1) the respiratory exchange ratio (R) exceeded 1.10, 2) achievement of 90% of age-predicted peak HR (220 − age), and 3) an RPE of 18 or more (Howley, Bassett, and Welch 1995; Midgley et al. 2007).

### 2.6 Psychological measurements

The participants’ RPE (Borg 1970) was recorded before and after the exercise intervention to assess psychological exercise intensity. Additionally, two-dimensional mood scale (TDMS) questionnaires were administered to assess psychological indicators before and after the exercise intervention. The TDMS is a momentary mood scale comprising two words describing the arousal and pleasure states (lively and relaxed) (Sakairi, Nakatsuka, and Shimizu 2013). Participants were asked to indicate how they felt about each mood-expressing word using an 11-point Likert scale ranging from –5 (listless) to 5 (lively) and –5 (irritated) to 5 (relaxed). In addition to “words” and “numbers” describing the psychological state, the shortened version (Ochi, Kuwamizu, Fujimoto, et al. 2022; Kuwamizu et al. 2022) used “person illustrations” and “color images” to reduce the burden of answering for participants who were unfamiliar with the experiment. The vitality level, which represents low arousal-displeasure to high arousal-pleasure, and stability level, which represents high arousal-displeasure to low arousal-pleasure, were measured. Based on these scores, pleasure level (vitality + stability) and arousal level (vitality − stability) were calculated.

### 2.7 fMRI measurements

All structural and functional brain images were acquired using a 3 T MRI scanner (Canon Medical Systems, Tochigi, Japan) with a 16-channel head coil. Anatomical images were acquired using a T1-weighted 3D magnetization-prepared rapid gradient echo sequence with the following parameters: inversion time = 900 ms, repetition time = 5.8 ms, echo time = 2.7 ms, flip angle = 9◦, slice thickness = 1.2 mm, field of view = 23×23 cm^2^, scan matrix = 256×256, number of slices = 160, and slice gap = non-gap. The fMRI images were acquired using ascending-order T2*-weighted gradient echo-planar imaging (EPI). The fMRI imaging conditions were as follows: repetition time, 2,000 ms; echo time, 25 ms; flip angle, 85◦; matrix, 64 × 64; effective field of view, 24 × 24 mm; and slice thickness, 3 mm to cover the whole brain.

### 2.8 fMRI data analysis

We performed image preprocessing and statistical analysis using Statistical Parametric Mapping (SPM12) revision 7487 (Wellcome Centre for Human Neuroimaging, London, UK) implemented in MATLAB 2023a (Mathworks, Natick, MA, USA). Functional images were realigned, slice timing corrected, and normalized to the MNI template (ICBM 152) with interpolation to a 2 × 2 × 2 mm space. The data were spatially smoothed (full width, half maximum [FWHM] = 8 mm) for univariate parametric modulation analysis. Motion and susceptibility artifacts were detected using the Art Toolbox (http://web.mit.edu/swg/software.htm). Outlier scans (head motion above 2 mm and/or changes in mean signal intensity above 4) identified by this procedure were then added as regressors of no interest for subsequent analyses. No participant was excluded after performing this quality check. To visualize the imaging results, the MRIcron software (https://people.cas.sc.edu/rorden/mricro/index.html) was used after modification.

### 2.9 Statistical analysis

All analyses were performed using R (4.3.2) and Rstudio (2023.06.0+421) software and the R package “anovakun.” Statistical significance was set at *P* < .05 for all comparisons. Mauchly’s sphericity test was used to determine whether sphericity was maintained. When a significant difference was observed, we conducted a repeated-measures two-way ANOVA with Greenhouse-Geisser’s epsilon correction. Otherwise, we conducted a repeated-measures two-way ANOVA. Significant differences obtained from two-way ANOVA were tested using the corresponding t-test with Shaffer’s modified sequentially rejective Bonferroni procedure. One-way ANOVA was performed on the pre- and post-exercise changes in reaction time for the N-back task, and a t-test with Shaffer’s modified sequentially rejective Bonferroni procedure was performed when a significant main effect was observed. As exploratory analyses, correlations among N-back performance, DLPFC, LC activity, and psychology parameters were examined using repeated measures correlation (R package “rmcorr”) (Bakdash and Marusich 2017). The rmcorr correlation coefficient (rrm) determines the common intra-individual relationship for paired measurements assessed on two or more occasions for multiple individuals (Barr et al. 2013).

We employed a summary statistics approach to delineate the neural substrates of task-related brain activity. In individual analyses, a general linear model was fitted to the fMRI data of each participant. Neural activity was modeled using delta functions convolved with a canonical hemodynamic response function. Task-related regressors for the RS, EX, and VR were implemented as regressors of interest. To control slow frequency fluctuations, a high-pass filter (256 s) was applied. Single-subject design matrices included six motion regressors and censored volumes as regressors, specified as nuisance regressors. Global scaling was applied. Parameter estimates from individual analyses comprised contrast images used for group-level analysis. The resulting voxel values for each contrast constituted a statistical parametric map of the *t* statistic (SPM{*t*}). The statistical threshold was set at *P* < .05 with family-wise error (FWE) correction at the cluster level for the entire brain, with a height threshold of *P* < .001. Anatomical locations were determined using the Atlas of the Human Brain, 4th edition, for anatomical labeling (Mai et al., 2015). To visualize the imaging results, we utilized MRIcron software (https://people.cas.sc.edu/rorden/mricro/index.html) with modifications.

Subsequently, a correlation analysis was conducted between 3-back performance and brain activity, focusing on brain structures known to influence N-back performance and mood, specifically the DLPFC and LC. Regions of interest (ROIs) were anatomically defined using the automated anatomical labeling atlas 3 (AAL3) (Rolls et al., 2020), and beta values were extracted from relevant ROIs.

## 3. Results

### 3.1 Overview

All participants completed the experiment without any reported adverse effects related to VR, such as motion sickness, dizziness, or headaches, after the VR condition.

### 3.2 Physiological and psychological parameter

HR and RPE were subjected to repeated-measures two-way ANOVA with condition (RS, EX, and VR) and session (before exercise and during/after exercise) as within-subject factors. A significant interaction between condition and session was observed for HR (*F*(2, 44) = 458.74, *P* < .001, *η^2^_p_* = 0.95) and RPE (*F*(1.61, 35.48) = 80.24, *P* < .001, *η^2^_p_* = 0.78). Regarding HR, the pre- and post-exercise changes showed a significant main effect of condition (*F*(2,44) = 458.74, *P* < .001, *η^2^_p_* = 0.95), with significantly higher values found in the EX (pre: 72.5 ± 5.7; post: 126.1 ± 10.5) and VR (pre: 71.7 ± 6.9; post: 124.0 ± 12.5) condition than in RS (pre: 75.2 ± 8.3; post: 75.3 ± 6.4) condition (EX: *t*(22) = 27.13, *P* < .001; VR: *t*(22) = 21.89, *P* < .001). Regarding RPE, the pre- and post-exercise changes showed a significant main effect of condition (*F*(1.61,35.48) = 80.24, *P* < .001, *η^2^_p_* = 0.78), with significantly higher values found in the EX (pre: 6.0 ± 0.0; post: 11.5 ± 2.4) and VR (pre: 6.0 ± 0.0; post: 11.2 ± 2.8) condition than in RS (pre: 6.0 ± 0.0; post: 6.0 ± 0.0) condition (EX: *t*(22) = 11.17, *P* < .001; VR: *t*(22) = 8.95, *P* < .001). No significant differences were found between the EX and VR conditions for HR and RPE, confirming that the experiments were conducted with comparable exercise loads. The increases in HR and RPE were comparable to those in a previous study where 10 min of moderate-intensity exercise was imposed (Ochi et al. 2018), suggesting that each participant performed the exercise at moderate intensity in this study.

Table 2 summarizes the results of the psychological mood states. Psychological mood states (vitality, stability, arousal, and pleasure) were measured using TDMS. A significant interaction between condition and session was observed for vitality (*F*(2, 44) = 6.53, *P* < .005, *η^2^_p_* = 0.23) and arousal level (*F*(2, 44) = 6.21, *P* < .005, *η^2^_p_* = 0.22). Regarding vitality level, the pre- and post-exercise changes showed a significant main effect of condition (*F*(2,44) = 6.53, *P* < .005, *η^2^_p_* = 0.23), with significantly higher values observed in the VR condition than in RS and EX conditions (vs. RS: *t*(22) = 4.08, *P* < .001; vs. EX: *t*(22) = 2.70, *P* < .001). Regarding arousal level, the pre- and post-exercise changes showed a significant main effect of condition (*F*(2,44) = 6.21, *P* < .001, *η^2^_p_* = 0.22), with significantly higher values found in the VR condition than in RS condition (*t*(22) = 3.55, *P* < .001). Regarding stability level, a significant main effect of the session was observed (*F*(1,22) = 5.73, *P* < .05, *η^2^_p_* = 0.20). No significant interactions or main effects were found at the pleasure level.

**Table 2.** TDMS during rest (RS), exercise (EX), and exercise under virtual reality (VR) conditions.

|  | RS |  | EX |  | VR |  |
| --- | --- | --- | --- | --- | --- | --- |
| Characteristic | Pre | Post | Pre | Post | Pre | Post |
| Vitality (point) | 0.5 (1.7) | 0.5 (1.6) | 1.0 (1.7) | 1.3 (1.8) | 0.5 (1.4) | 1.9 (1.7)*† |
| Stability (point) | 1.5 (1.9) | 1.5 (2.0) | 1.4 (1.8) | 0.6 (1.9) | 1.4 (1.8) | 0.6 (1.8) |
| Pleasure (point) | 2.0 (3.3) | 2.0 (3.3) | 2.3 (3.3) | 1.9 (3.1) | 1.9 (2.8) | 2.5 (3.0) |
| Arousal (point) | -1.0 (1.5) | -1.0 (1.6) | -0.4 (1.2) | 0.7 (2.0) | -0.9 (1.5) | 1.3 (1.7)* |

Mean (SD).
Note: RS=rest without the head-mounted display condition; EX=exercise without head-mounted display condition; VR=exercise with a head-mounted display condition; SD=standard deviation; TDMS=two-dimensional mood scale.
\* vs. RS condition.
† vs. EX condition.

### 3.3 Working memory performance: N-Back task

First, for each condition, we examined each task to determine whether any changes were observed before and after the exercise. In the 0-back task, a main effect of time was observed (*F*(1,22) = 12.6419, *P* < .05, *η^2^_p_* = .36), with a reduction in reaction time after exercise (Fig. 3A). In the 1- and 3-back task, an interaction between time and condition was noted (1-back: *F*(2,44) = 3.2993, *P* < .05, *η^2^_p_* = .13, Fig. 3B; 3-back: *F*(2,44) = 5.6327, *P* < .05, *η^2^_p_* = .21, Fig. 3C). No significant differences were found in the Post-hoc test in the 1-Back task; however, in the 3-back task, the paired t-test with Shaffer’s modified sequentially rejective Bonferroni procedure showed significant differences between the rest and VR conditions (*t* = 2.3399, *P* < .05) and between the EX and VR conditions after exercise (*t* = 2.7109, *P* < .05). Subsequently, in the 1- and 3-back tasks, we checked the amount of change before and after the exercise (post -pre session) to ascertain any differences between conditions. In the 1-back task, a main effect across conditions was evident (*F*(2,44) = 3.2993, *P* < .05, *η^2^_p_* = .13), with a significant difference trend between RS and VR (*t* = 2.53, *P* = .057, Fig. 3D). In the 3-Back task, a main effect across conditions was observed (*F*(2,44) = 5.6327, *P* < .05, *η^2^_p_* = .20), with significant differences between RS and VR (*t* = 3.7703, *P* < .005) and between EX and VR (*t* = 2.6577, *P* < .05) (Fig. 3E).

No significant main or interaction effects were observed for the 0- and 1-back tasks regarding the task correctness. A significant main effect of time was found for the 3-back task (*F*(1,22) = 4.3839, *P* < .05, *η^2^_p_* = .17), confirming that the percentage of correct responses increased before and after the exercise; however, no differences were observed between conditions (Table 3).

**Table 3.** Accuracy of N-back task under rest (RS), exercise (EX), and e exercise with a head-mounted display condition (VR)

|  | RS |  | EX |  | VR |  |
| --- | --- | --- | --- | --- | --- | --- |
| Characteristic | Pre | Post | Pre | Post | Pre | Post |
| 0-back (%) | 99.8 (0.8) | 99.6 (1.3) | 100.0 (0.0) | 99.4 (2.4) | 99.8 (0.8) | 99.8 (0.8) |
| 1-back (%) | 99.0 (2.2) | 98.8 (2.2) | 98.9 (1.8) | 98.7 (1.8) | 97.9 (3.3) | 99.0 (2.3) |
| 3-back (%) | 81.9 (8.2) | 84.5 (8.2) | 78.7 (9.4) | 83.1 (9.0) | 84.3 (7.9) | 84.5 (8.6) |
| Mean (SD). |  |  |  |  |  |  |

Note: RS=rest without the head-mounted display condition; EX=exercise without head-mounted display condition; VR=exercise with a head-mounted display condition; SD=standard deviation.

**Fig. 3.**
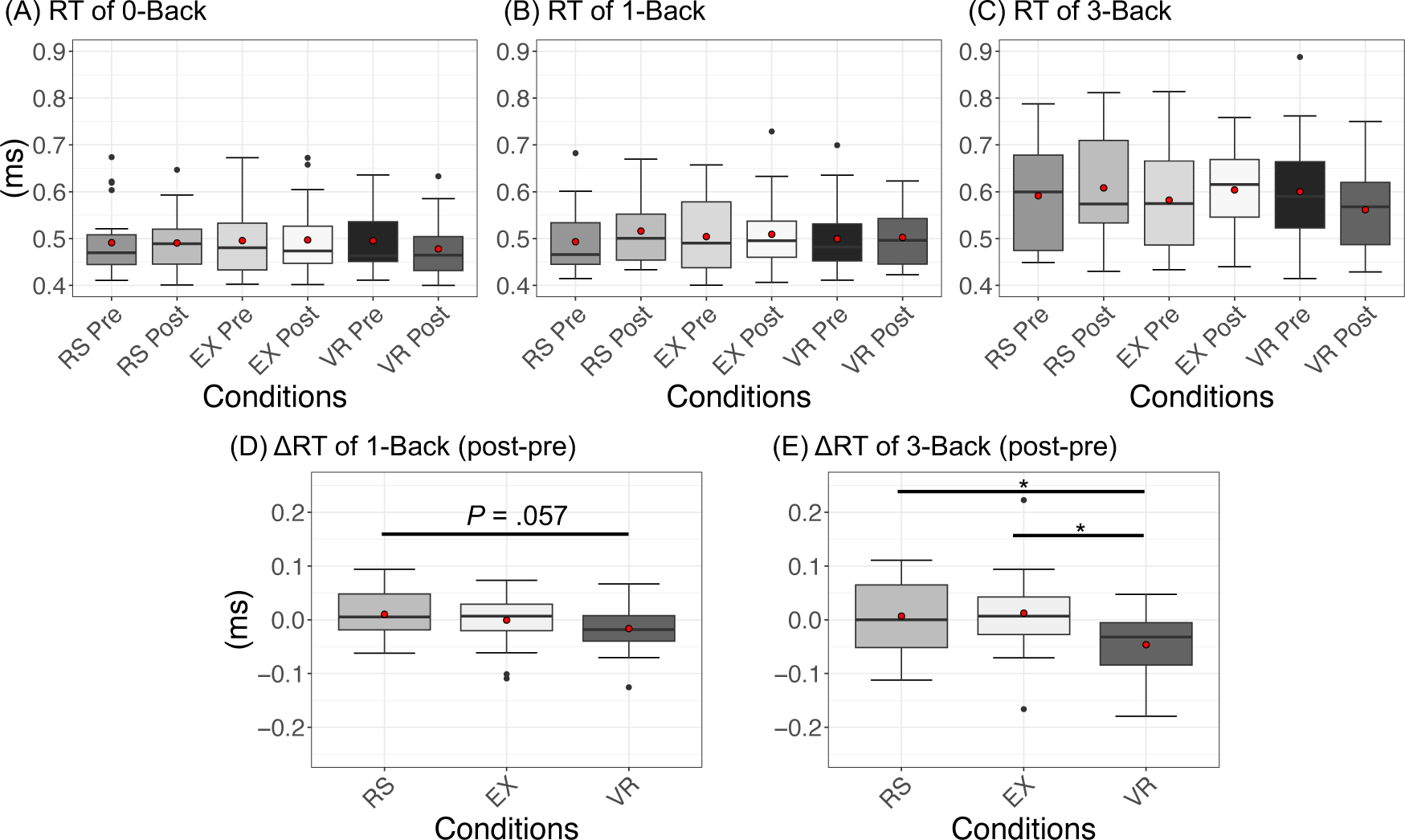
RT for the 0-(A), 1-(B), and 3-back tasks (C). Changes in RT between pre- and post-sessions for the 1-(D) and 3-back tasks (E). A significant difference trend was found between RS and VR in the 1-back task (*P* = .057). Significant differences were found between RS and VR and between EX and VR in the 3-back task (*P* < .05). The upper and lower ends of the whiskers represent the highest data points within 1.5 IQRs of the upper quartiles and the lowest data points within 1.5 IQRs of the lower quartiles, respectively. The bands inside the boxes indicate the medians. The red circle is the mean. RS=rest condition; EX=exercise condition; RT=reaction time; VR=exercise with a head-mounted display condition; IQR=interquartile range.

### 3.4 fMRI results

The results of the fMRI results indicated that the supplementary motor cortex, inferior parietal gyrus, and left precentral gyrus were activated during the 3-back task 10 min before the activity (Pre). In contrast, during the 3-back task after 10 min of activity (Post), the following regions were activated: supplementary motor area, inferior parietal gyrus, and left precentral gyrus in RS. Moreover, in EX and VR conditions, activation was observed in the supplementary motor area, inferior parietal gyrus, left precentral gyrus, and right superior frontal gyrus. The brain regions activated in the EX and VR overlapped in many areas. However, brain regions activated in VR, specifically the left insula and left DLPFC, did not show activation in EX (Table 4 and Fig. 4).

**Fig. 4.**
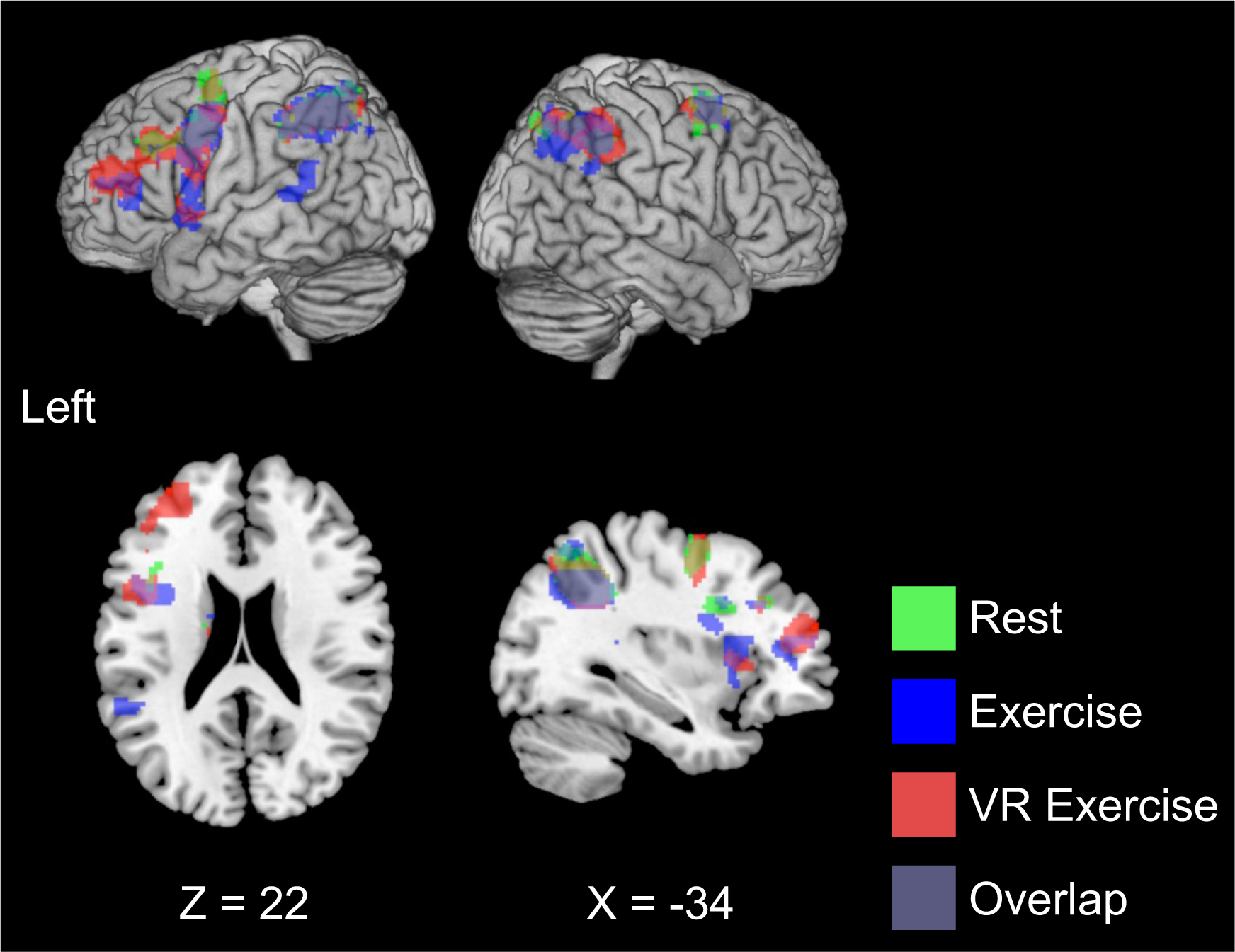
Brain activity during the 3-back task after 10 min of rest (RS) (green region), exercise (EX) (blue region), and exercise with a head-mounted display condition (VR) (red region)

**Table 4.**
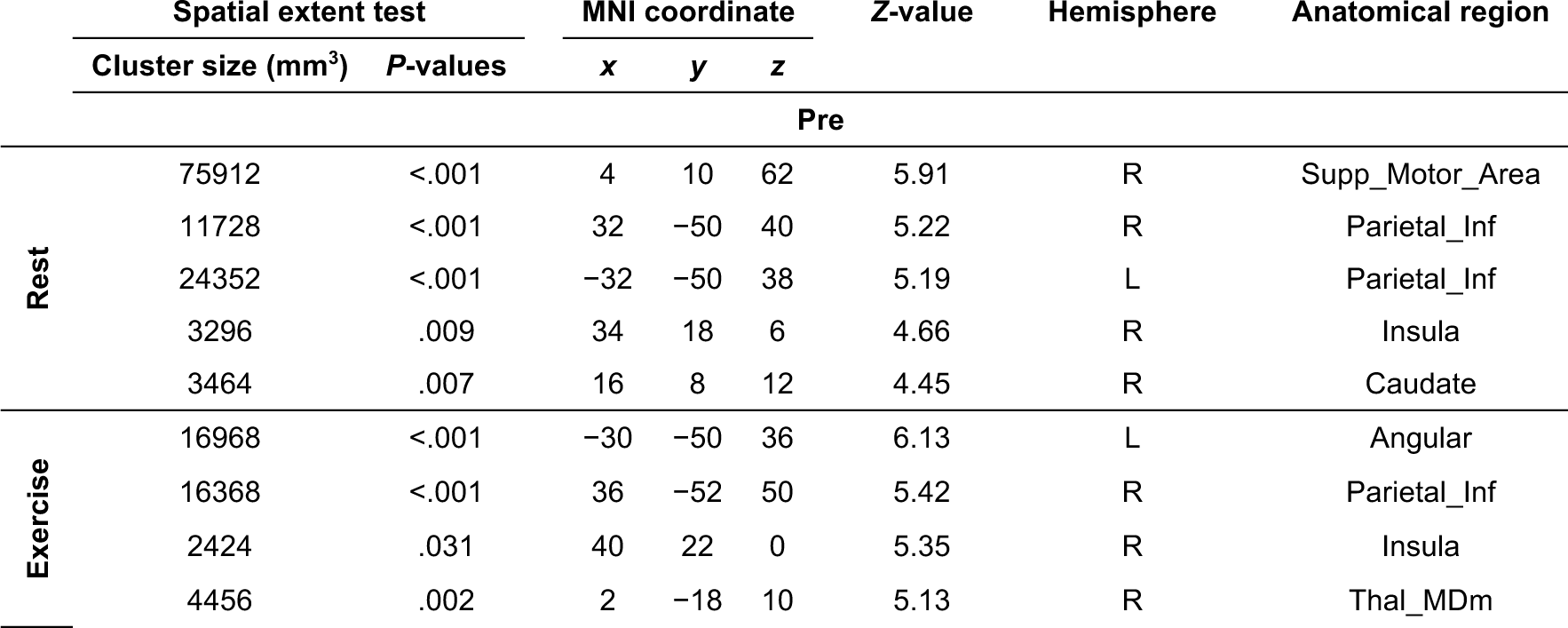
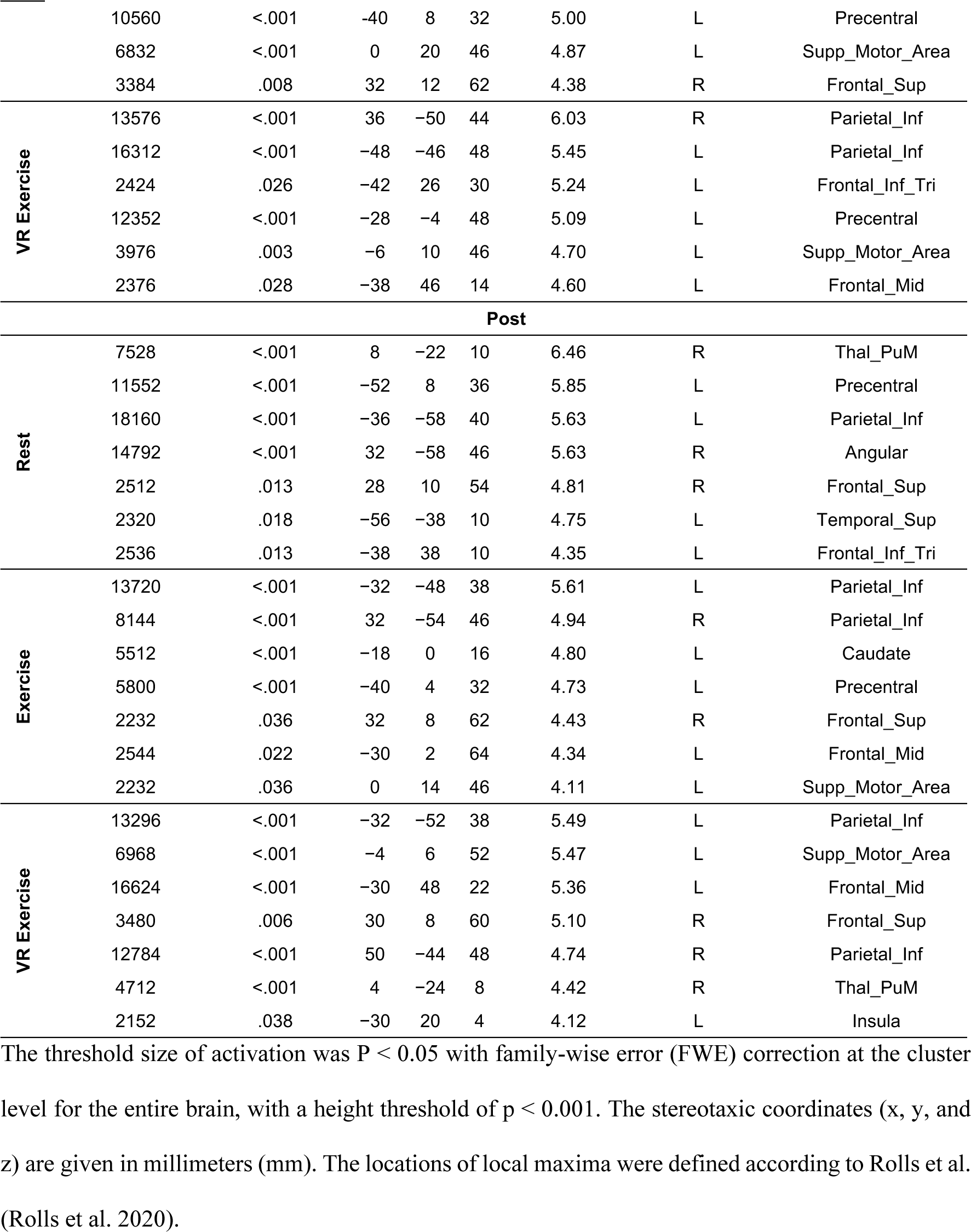
Significant clusters of brain activity in the 3-back task before and after 10 min of RS, EX, and VR.

### 3.5 Association of N-back performance and fMRI results

First, we investigated the brain regions across the entire brain about N-back task performance. The results indicated that no brain regions were associated with changes in the reaction time in the 3-back task, which varied before and after the VR exercise.

Table 2 presents the results of the repeated measures correlation analyses. In this study, we specifically focused on the activity of the DLPFC and LC, which we hypothesized would be associated with increased N-back performance and mood. However, we found no significant correlation between reaction time changes in the 3-back task and activity in the right and left DLPFC and the right and left LC.

Furthermore, we examined the psychological parameters related to DLPFC and LC activity during the 3-back task. No significant correlations were found with DLPFC and LC for all mood indices (Table 5).

**Table 5.** Repeated measures correlation analysis for DLPFC, LC activity, reaction time for the 3-back task, and psychology parameters.

|  |  | Right DLPFC | Left DLPFC | Right LC | Left LC |
| --- | --- | --- | --- | --- | --- |
| RT of 3-back | *r* | .007 | -.040 | .028 | .021 |
|  | *P* | .94 | .66 | .77 | .82, |
|  | 95% CI | -.176-.189 | -.221-.143 | -.155-.209 | -.162-.203 |
| Vitality | *r* | .037 | .094 | -.090 | -.103 |
|  | *P* | .69 | .32 | .34 | .27 |
|  | 95% CI | -.146-.218 | -.090-.271 | -.268-.094 | -.281-.080 |
| Stability | *r* | -.021 | -.041 | .075 | .081 |
|  | *P* | .82 | .66 | .42 | .39 |
|  | 95% CI | -.202-.162 | -.222-.142 | -.109-.254 | -.103-.260 |
| Pleasure | *r* | .010 | .033 | -.007 | -.012 |
|  | *P* | .92 | .73 | .94 | .90 |
|  | 95% CI | -.173-.192 | -.151-.214 | -.189-.176 | -.194-.171 |
| Arousal | *R* | .044 | .101 | -.124 | -.139 |
|  | *P* | .64 | .28 | .19 | .14 |
|  | 95% CI | -.140-.224 | -.083-.278 | -.300-.060 | -.313-.045 |

Note: RT=reaction time; DLPFC=the dorsolateral prefrontal cortex; LC=locus coeruleus.

### 3.6 Association of psychological parameter and N-Back performance

Finally, the psychological parameters related to N-back performance were examined. The vitality of TDMS was negatively correlated with the 3-back reaction time (*r* = −.224, *P* < .05, 95% confidence interval [CI] = −.390–−.043; Fig. 5). Stability (*r* = −.005, *P* = .96, 95% CI = −.187–.178), pleasure (*r* = −.149, *P* = .11, 95% CI = −.323–.034), and arousal (*r* = −.160, *P* = .09, 95% CI = −.333–.023) of TDMS were not significantly related.

**Fig. 5.**
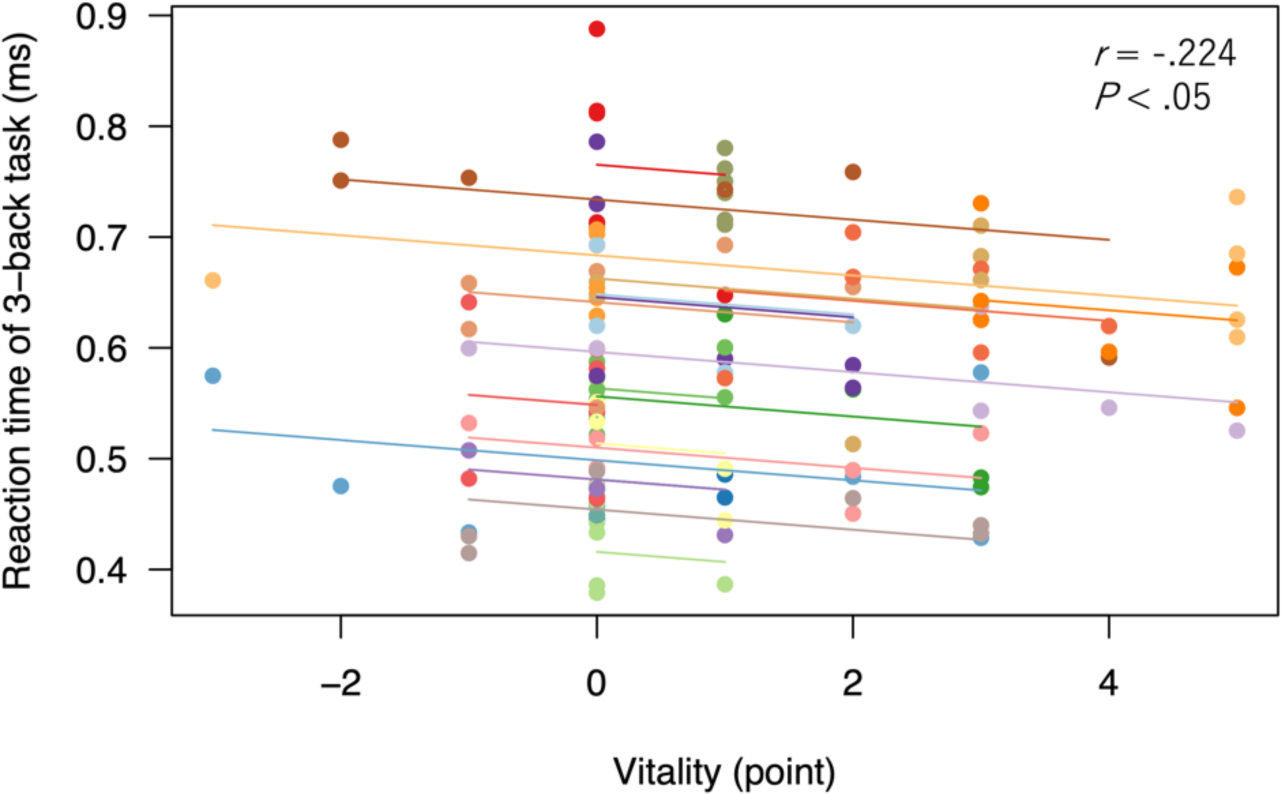
Repeated measures correlation analysis for pre- and post-exercise vitality and reaction time for the 3-back task in each condition. The reaction time to the 3-back task decreased with increasing vitality before and after exercise and rest in the three conditions. Results from the same participant were given the same color, with corresponding lines to show the rmcorr fit for each participant.

## 4. Discussion

In this study, we used fMRI to investigate whether VR exercise could improve working memory function and mood and to determine the involvement of the DLPFC and LC in this cognitive mechanism. The results showed that VR exercise increased vitality and improved working memory function. Notably, we found that reaction times in the 3-back task decreased as vitality increased. However, no direct relationship was observed between the enhancement of working memory induced by VR exercise and changes in DLPFC and LC activity during working memory task post-exercise. This study suggests that mood enhancement plays a role in improving working memory function and that VR exercise can be an effective method to achieve this.

### 4.1. VR exercise enhances working memory function

HR and RPE, which reflect exercise intensity, were increased to the same extent as in previous studies that used 10-min moderate-intensity exercise interventions (Yanagisawa et al. 2010; Ochi et al. 2018; Suwabe et al. 2021, 2017). These results suggest that the EX and VR conditions in this study can be considered moderate-intensity exercises. Because no differences were observed in the HR between the conditions, the EX and VR conditions induced the same exercise intensity.

We examined the impact of exercise on executive function. The results showed that in the 3-back task, the reaction time was reduced in the VR condition compared to the RS and EX conditions. These results indicate that exercise under VR improves executive function compared to rest and exercise alone. However, the 3-back reaction time in the EX condition was unchanged from the RS condition and did not induce any improvement in executive function. A previous study reported that 10 min of moderate-intensity bicycle exercise did not improve working memory function (Yamazaki et al. 2018), and the EX condition in this study replicated this result. In this study, the effect of improving executive function after exercise was observed in the VR condition compared to the EX condition, suggesting that the environment in which exercise is performed and the exercise itself may be important for improving executive function.

### 4.2. VR exercise enhances mood

Furthermore, we examined the effects of VR exercise on mood. Post-exercise activation was higher in the VR condition than in the rest (RS) and traditional exercise (EX) conditions and both vitality and arousal were higher in the VR condition than in the RS condition. We previously reported that 10 min of VR exergaming increases vitality and arousal (Ochi, Kuwamizu, Fujimoto, et al. 2022). The results of this study showed that the VR exercise replicated these mood-enhancing effects. Furthermore, vitality was negatively correlated with reaction time in the 3-back task, indicating that as vitality increased, reaction time decreased. These results are consistent with previous studies (Suwabe et al. 2021), which reported that combining music with exercise may enhance executive function. These findings suggest that working memory function improves as vitality increases with exercise and that VR exercise interventions specifically enhance this effect. In this study, an increase in vitality was observed only in the VR exercise condition, indicating that VR exercise is a beneficial program that enhances both mood and working memory.

### 4.3. Brain regions during the N-Back task

In this study, we used fMRI to investigate the brain regions activated during the N-back task. The brain areas activated during the 3-back task included the right medial pulvinar, left precentral gyrus, left inferior parietal gyrus, right angular gyrus, right superior frontal gyrus, left superior temporal gyrus, and left inferior frontal gyrus. These activated regions were also included in the regions from a meta-analysis evaluating active brain regions during the N-back task (Z. A. Yaple, Stevens, and Arsalidou 2019). In all conditions, no areas showed significantly increased or decreased activity before or after exercise. However, activation of the left supplementary motor area, left insula, and left DLPFC was observed after exercise in the VR condition, coinciding with improved performance in the 3-Back task. DLPFC activity, crucial in N-back task performance (Owen et al. 2005), increases with increasing difficulty (Lamichhane et al. 2020). Therefore, we hypothesized that increased activity in the left DLPFC is involved in improving 3-back task performance and proceeded with the analysis.

### 4.4. Brain regions involved in improving working memory function through VR exercise

We investigated brain regions associated with improved executive function under VR motor conditions. Our whole-brain analysis did not identify any regions correlated with shorter reaction times for the 3-back task in the VR motor condition. Subsequently, we directed our analysis toward the hypothesized relationship between the DLPFC and LC activity and changes in executive function following VR exercise. Nonetheless, no association was found between activity in the left or right DLPFC and LC and shorter reaction times in the 3-back task. Prior research using tasks such as the color-word Stroop task has suggested that increased DLPFC activity is linked to exercise-induced improvements in executive function (Yanagisawa et al. 2010; Byun et al. 2014; Hyodo et al. 2012; Damrongthai et al. 2021). Given the DLPFC’s role in both N-back and color-word Stroop tasks (Lamichhane et al. 2020), we postulated that enhanced DLPFC activity would improve performance in the 3-back task with VR exercise. Contrary to these precedents, our study did not find a relationship between exercise-induced improvement in 3-back task performance and increased DLPFC activity.

Previous studies have demonstrated that pupil dilation during exercise (which may be related to LC activity) predicts the effect of improved Stroop task performance (Kuwamizu et al. 2022; 2023, Yamazaki et al. 2023). The present study found no relationship between LC activity and working memory function enhancement, possibly because LC activity was measured during the post-exercise working memory task rather than during exercise itself. Several pupillary studies have also reported that the cognitive enhancement effect of exercise is not related to pupil size during post-exercise cognitive tasks (Shigeta et al. 2021; McGowan et al. 2019), consistent with our findings regarding LC activity post-exercise. LC activity during VR exercise may be related to the working memory enhancement effect induced by VR. In our study, the impact of VR exercise on pupil diameter remained unclear as HMDs capable of measuring pupil diameter were not utilized. Future research measuring pupil diameter during VR exercises may shed light on how increased brain arousal through LC activity serves as a neural mechanism for enhanced working memory function.

Although specific regional activity is critical, the connectivity between these regions also plays a vital role in cognitive task performance (Yu and Liu 2021). The cognitive task in this study was too brief in the resting state to allow for the assessment of neural connectivity; however, future studies should employ task paradigms designed to evaluate this aspect.

### 4.5. Limitations and perspectives

Our sample size was comparable to that of previous studies that used functional brain imaging (Suwabe et al. 2017, 2018; Ochi et al. 2018; Ochi, Kuwamizu, Suwabe, et al. 2022; Suwabe et al. 2021). However, further validation assuming diverse individual differences is required to elucidate the mechanisms of the effects of VR exercise on working memory functions. Furthermore, the study’s emphasis on healthy young adults raises the potential for replicating similar findings among older individuals and those with mental health conditions. These insights underscore VR’s potential as a novel exercise modality with benefits extending beyond exercise adherence to potentially preventing dementia and depression. Finally, the participants in this study had no experience with VR, raising uncertainties about whether factors such as habituation or boredom from prolonged VR intervention may impact working memory enhancement. Future verification of the long-term intervention effects of VR exercises will enable us to propose a new exercise program using VR.

## 5. Conclusions

This study demonstrates that VR exercise improves mood and working memory function. Although the precise neural mechanisms underlying these effects remain unclear, our findings suggest that an enhanced state of high arousal and pleasure mood is crucial for improving working memory function, with VR being a potent motor factor in achieving this condition. To encourage the adoption of VR exercise for habitual exercise and the enhancement of mood and executive function, further exploration into the mechanisms underlying the improvement of executive function through VR exercise is warranted. Additionally, validating whether these effects can be reproduced in distinct populations such as children and older adults is essential.

## Data availability statement

The datasets generated and/or analyzed during this study are not publicly available but are available from the corresponding author upon reasonable request.

## Acknowledgments

We would like to thank all participants of the study and Editage (www.editage.com) for English language editing.

